# A single *Verticillium dahliae* effector determines pathogenicity on tomato by targeting auxin response factors

**DOI:** 10.1101/2022.11.22.517554

**Authors:** Jinling Li, Luigi Faino, Gabriel L. Fiorin, Sagar Bashyal, Arno Schaveling, Grardy C.M. van den Berg, Michael F. Seidl, Bart P.H.J. Thomma

## Abstract

**SUMMARY:** *Verticillium dahliae* is a xylem-invading fungal pathogen that causes devastating vascular wilt diseases on hundreds of plant hosts, including tomato (*Solanum lycopersicum*). Although individual *V. dahliae* strains are typically characterized by their broad host range, differential pathogenicity occurs on nearly all hosts. Currently, the molecular basis underlying such pathogenicity differences remains unknown. We used comparative genomics to identify a single effector gene that specifically occurs in tomato-pathogenic *V. dahliae* strains and is expressed during tomato colonization. Functional analyses showed that this Tom1 effector governs pathogenicity on tomato, as *Tom1* deletion prohibited tomato colonization, while introduction of *Tom1* into non-pathogenic *V. dahliae* or into saprophytic sister species *V. tricorpus* and *V. nubilum* resulted in disease. Through proteomics-based approaches, auxin response factors (ARFs) were identified as *in planta* targets of Tom1. Intriguingly, repression of *SlARF2a* expression by virus-induced gene silencing fully impaired *V. dahliae* colonization of tomato, solidifying its role as susceptibility target.Collectively, our findings indicate that a single effector, Tom1, mediates pathogenicity of *V. dahliae* on tomato by targeting auxin response factors.

## INTRODUCTION

In their co-evolutionary arms race with plant hosts, adapted pathogens evolved molecules to counteract host immunity or interfere with host physiology to support pathogen colonization (Cook et al., 2015; Lo Presti *et al*., 2015; Ngou et al., 2022). This host manipulation is mediated by effector proteins, which typically are small, cysteine-rich that are specifically secreted by pathogens during plant colonization (Rovenich *et al*., 2014; De Wit, 2016; Wilson and McDowell, 2022). Accordingly, the identification and functional characterization of effectors is important for mechanistic understanding of microbial pathogenesis to ultimately provide valuable knowledge to develop effective disease management strategies (Rovenich *et al*., 2014). It is generally observed that effector activities are redundant, and that single effectors are dispensable for full pathogen virulence. Thus, effectors mostly contribute quantitatively to the host colonization ability of a pathogen (de Jonge *et al*., 2011; Win *et al*., 2012; Tan *et al*., 2015; Wilson and McDowell, 2022). Nevertheless, some effectors are fully responsible for the ability to cause disease as so-called pathogenicity determinants. For instance, the PtrToxA effector from the wheat tan spot pathogen *Pyrenophora tritici-repentis* confers the ability to cause disease on wheat cultivars carrying the corresponding susceptibility gene Tsn1 (Friesen *et al*., 2006). Expression of PtrToxA in a nonpathogenic *P. tritici-repentis* isolate renders the isolate pathogenic, indicating that the effector is sufficient for *P. tritici-repentis* pathogenicity in wheat (Ciuffetti *et al*., 1997).

*Verticillium* is a genus of Ascomycete fungi consisting of ten soil-borne, asexual species with lifestyles ranging from saprophytic to pathogenic (Klosterman et al., 2009; Inderbitzin and Subbarao, 2014; Klimes et al., 2015). Among these species, *V*. *isaacii, V. klebahnii, V. zaregamsianum, V. tricorpus* and *V*. *nubilum* are saprophytes that occasionally cause opportunistic infections on plants that have been weakened by other stresses (Inderbitzin et al., 2011; Gurung et al., 2015; Seidl et al., 2015; Shi-Kunne et al., 2018). In contrast, *V. dahliae, V. albo-atrum, V. alfalfae, V. nonalfalfae* and *V. longisporum* are genuine pathogens that cause vascular wilt diseases, although their host ranges differ substantially (Pegg and Brady, 2002; Fradin and Thomma, 2006; Inderbitzin et al., 2011; Depotter et al., 2016). For example, whereas *V*. *longisporum* causes disease on brassicaceous hosts such as rapeseed and cauliflower (Depotter et al., 2016), and *V. alfalfae* only infects lucerne (Inderbitzin et al., 2011), *V. dahliae* is the most notorious pathogen of the genus that can infect hundreds of species (Fradin and Thomma, 2006). Interestingly, individual *V. dahliae* strains display differential pathogenicity among hosts (Schnathorst & Mathre, 1966; Schnathorst & Sibbett, 1971).

Tomato (*Solanum lycopersicum*) is one of the most valuable vegetable crops worldwide and an important host of *V. dahliae*. The first source of genetic *V. dahliae* resistance was found in the tomato accession Peru Wild to encode the immune receptor Ve1 (Schaible *et al*., 1951; Kawchuk et al., 2001; Fradin et al., 2009). Whereas race 1 strains are contained by *Ve1*, resistance-breaking race 2 isolates are ubiquitous (Dobinson et al., 1996). Based on comparative genomics, the Ave1 effector that is recognized by Ve1 was identified, revealing that race 2 strains overcame Ve1-mediated recognition by purging of *Ave1* (de Jonge et al., 2012; Faino et al., 2016). Importantly, Ave1 promotes *V. dahliae* virulence on plants that lack *Ve1* through host microbiota manipulation through its selective antimicrobial activity targeted against antagonistic bacteria of the Sphingomonadales order (Snelders *et al*., 2020). Intriguingly, a multiallelic *Ave1* homolog was identified as *VdAve1-like* (*VdAve1L*), the products of which are not recognized by Ve1, and of which the VdAve1L2 variant promotes virulence on tomato through targeting of antagonistic Actinobacteria (Snelders et al., 2022).

Recently, the *SlV2* locus of *Solanum neorickii* was found to provide resistance against *V. dahliae* race 2 strains, although resistance-breaking race 3 strains were reported simultaneously (Usami *et al*., 2017). Based on comparative population genomics, the *V. dahliae* Av2 effector was identified as the corresponding avirulence molecule (Chavarro-Carrero *et al*., 2021). Interestingly, whereas no *Ave1* allelic variation was found, and race 2 strains escape recognition solely through *Ave1* purging, two allelic *Av2* variants were found that encode Av2 variants that differ by a single amino acid, although both activate *Av2*-mediated immunity (Chavarro-Carrero *et al*., 2021). Accordingly, resistance-breaking race 3 strains all lack the *Av2* gene (Chavarro-Carrero *et al*., 2021). Thus far, a role of Av2 in virulence could not be demonstrated and its molecular function remains enigmatic (Chavarro-Carrero *et al*., 2021).

Intriguingly, not all *V. dahliae* strains cause disease on tomato, even in absence of resistance genes. Here, we sought to identify the molecular basis of the ability of *V. dahliae* strains to cause disease on tomato through comparative and functional genomics.

## MATERIALS AND METHODS

### Pathogenicity assays

*V. dahliae* inoculations were performed on two-week-old tomato cultivar Moneymaker, and outgrowth assays, canopy measurements and biomass quantifications were performed as described (Fradin et al., 2009; Santhanam et al., 2013; Song et al., 2018). Disease symptoms were scored up to 21 days post inoculation (dpi).

### Phylogenetic analysis

A *V. dahliae* phylogenetic tree was generated by REALPHY (version 1.12) (Bertels et al., 2014) using Bowtie2 (Langmead and Salzberg, 2012) to map reads against the *V. dahliae* strain JR2 genome assembly (Faino et al., 2015). A maximum likelihood phylogenetic tree was inferred using RAxML (version 8.2.8) with the GTRGAMMA model and 100 bootstrap replicates (Stamatakis, 2014). *V. alfalfae* strain ms102 was used to root the tree.

### Comparative genomics

Short reads were mapped onto the *V. dahliae* strain JR2 genome assembly (Faino et al., 2015) using BWA (Li and Durbin, 2010) with default options. R scripts were used to identify presence/absence polymorphism, and genomic regions were considered present if the coverage depth was ≥5x, while those with coverage depth <5x or with partial coverage were considered absent. Genomic regions that are only present in tomato-pathogenic strains were determined, and genes were extracted using an R script. SignalP (version 4.1) (Petersen et al., 2011) and EffectorP (version 1.0) (Sperschneider et al., 2016) were used to identify signal peptides and effector genes, respectively.

### Deletion and complementation strains

Sequences flanking the Tom1 coding sequence were amplified from genomic *V. dahliae* DNA using primers KO-Tom1-LBF and KO-Tom1-LBR, and primers KO-Tom1-RBF and KO-Tom1-RBR (Table S1), cloned into pRF-HU2 (Frandsen et al, 2008) and transformed into *V. dahliae* strain JR2 as described (Santhanam et al., 2013). Transformants were selected on PDA containing cefotaxime (200 μg/mL) and hygromycin B (50 μg/mL) (Duchefa, Haarlem, the Netherlands), and *Tom1* absence was verified with PCR.

The *Tom1* coding sequence plus 1.2 kb upstream and 1.1 kb downstream (*pTom1::Tom1*) was amplified using primers Tom1-com-F and Tom1-com-R (Table S1), cloned into the Gateway^™^ compatible vector PCG (Zhou et al., 2013) and transformed into the *Tom1* deletion mutant, *V. dahliae* strain ST100, *V. tricorpus* strain MUCL9792, and *V. nubilum* strain PD397. Transformants were selected on PDA supplemented with cefotaxime (200 μg/mL) and geneticin (25 μg/mL) (Sigma-Aldrich Chemie BV, Zwijndrecht, The Netherlands) and successful transformation was confirmed with PCR.

### Constructs for transient expression *in planta*

The nucleotide sequence encoding mature Tom1 without signal peptide was amplified by PCR using cDNA of tomato infected with *V. dahliae* strain JR2 as template. The nucleotide sequence encoding mature Sue1 without signal peptide was amplified by PCR using cDNA from cotton infected by *V. dahliae* strain CQ2. The nucleotide sequence encoding the signal peptide of the *Nicotiana tabacum PR1a* gene (PR1aSP) was amplified by PCR using PR1aSP as template. Amplicons were merged by means of overlapping-extension PCR and cloned into the Gateway entry vector pDONR207 (Invitrogen, Bleiswijk, the Netherlands). Expression clones were generated for production of GFP-fusion proteins (PR1aSP-Tom1::pSOL2095 and PR1aSP-Sue1::pSOL2095) or Myc-fusion proteins (PR1aSP-Tom1::pGWB20 and PR1aSP-Sue1::pGWB20). The nucleotide sequences encoding the SlARF1 and SlARF2a proteins were amplified by PCR using cDNA from tomato leaves. Amplicons were cloned into the Gateway vector pDONR207 (Invitrogen, Bleiswijk, the Netherlands) and expression clones were generated for production of GFP-fusion proteins (SlARF1::pSOL2095 and SlARF2a::pSOL2095). Primer sequences are in Table S1. All expression clones were verified by sequencing.

### Co-immunopurification

Electrocompetent *Agrobacterium tumefaciens* strain GV3101 cells were transformed with PR1aSP-Tom1::pSOL2095 or PR1aSP-Sue1::pSOL2095. Colonies were picked and grown overnight in YEB (5 g/L beef extract, 5 g/L bacteriological peptone, 5 g/L sucrose, 1 g/L yeast extract and 2 g/L MgSO_4_) supplemented with 50 μg/L of kanamycin and 25 μg/L of rifampicin at 28°C and 200 rpm. Cultures were pelleted for 8 minutes at 5.000 *x* g at room temperature and resuspended in MMA (1X Murashige and Skoog basal salt mixture, 10 mM MES, 20 g/L sucrose; 200 μM acetosyringone; pH 5.6). Subsequently, leaves of six-week-old *Nicotiana benthamiana* were transiently transformed by *A. tumefaciens-mediated* expression in quadruplicates with PR1aSP-Tom1::pSOL2095 or PR1aSP-Sue1::pSOL2095 constructs and the P19 silencing suppressor in 1:1 ratio at final optical density (OD_600_) of 0.8 for each construct. After three days, leaves were frozen in liquid nitrogen and ground to fine powder. Proteins were extracted using extraction buffer (150 mM NaCl, 1.0% [v:v] IGEPAL CA-630 [NP-40], 0.5% [w:v] sodium deoxycholate, 0.1% [w:v] SDS, 50 mM tris; pH 8.0), plus one EDTA-free protease inhibitor tablet (Roche, Mannheim, Germany) per 50 mL of extraction buffer, and 1 g of sample per 2 mL of extraction buffer was used. After centrifugation at 21.000 *x* g for 15 min at 4°C supernatants were transferred to new tubes. Next, 10 mL of protein extract was incubated with 60 μL (50% slurry) of GFP-trap magnetic agarose beads (ChromoTek, Planegg-Martinsried, Germany) for 3 h on a rotary shaker at 4°C. Next, beads were collected using a magnetic rack and washed three times in 1 mL extraction buffer. Next, beads were mixed with Laemmli buffer (Laemmli, 1970), boiled for 10 min and loaded onto SDS-PAGE (Bio-Rad, California, USA). A short electrophoresis run (5 minutes at 200 V) in TGS buffer (Bio-Rad, California, USA) was performed, followed by staining with Coomassie Brilliant Blue (Bio-Rad, California, USA). Single bands were cut using clean razor blades and stored in 1X PBS buffer (Thermo Fisher Scientific, California, USA) at 4°C. Further processing of the gel slices and LC-MS/MS runs (TripleTOF5600+ system) were performed at the Beijing Genome Institute (BGI, Hong Kong, China). For other co-immunopurifications, the same procedure was followed, except that 2 mL of total protein extract was incubated with 15 μL (50% slurry) of the GFP-trap magnetic agarose beads for 1 h on a rotary shaker at 4°C. Additionally, samples were subjected to electrophoresis in TGS buffer for 60 minutes at 200 V, and proteins were transferred to nitrocellulose membrane at 100 V for 60 minutes using transfer buffer (25 mM tris, 190 mM glycine, 20% methanol). Further steps were performed as described previously (Liebrand *et al*., 2012). *α*GFP-HRP antibody (Miltenyi Biotec GmbH, Bergisch Gladbach, Germany), αMyc antibody (Santa Cruz Biotechnology, Heidelberg, Germany) and αMouse-HRP antibody (GE Healthcare, Eindhoven, the Netherlands) were used.

### RESULTS

#### *Verticillium dahliae* pathogenicity on tomato

Previously, a suite of *V. dahliae* strains have been sequenced in our lab, of which ST100 lacks the ability to infect tomato (de Jonge et al., 2012, 2013; Faino et al., 2015; Chavarro-Carrero et al., 2021; Torres et al., 2021). To identify additional strains that are non-pathogenic on tomato, we generated a phylogenetic tree of all sequenced *V. dahliae* strains (Table S2) revealing the previously described three major lineages (Figure 1a) (Chavarro-Carrero *et al*., 2021; Torres et al. 2021). *V. dahliae* strains v781I, TM6, CQ2, V991, V117, V1381I, 463, V76 and T9 clustered with the tomato non-pathogenic strain ST100, while the remaining strains were found to be more related to previously characterized tomato-pathogenic strains (Figure 1a). Of particular interest were strains V574, V700, v679 and Vd39 that belong to the same sub-lineage as ST100 but are phylogenetically distinct (Figure 1a), and we therefore selected strains V991, V117, T9, V574 and Vd39 to evaluate their pathogenicity on tomato. In contrast to strain JR2, which induced clear stunting on tomato, all these strains failed to cause visible symptoms on tomato (Figure 1b,c). Plating stem of sections of the inoculated plants on agar medium confirmed absence of the fungus in the stems of inoculated tomato plants (Figure 1d). Collectively, these data demonstrate that stains ST100, V991, V117, T9, Vd39 and V574 are non-pathogenic on tomato.

**Figure 1.**
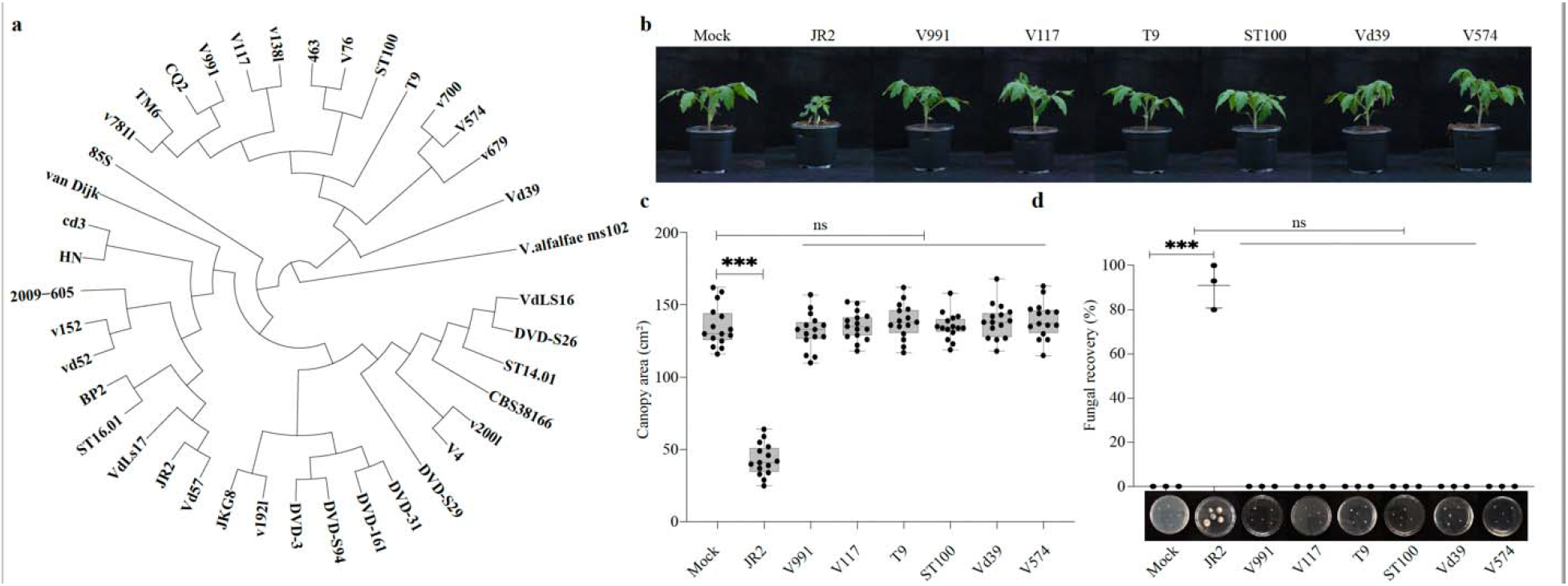
*Verticillium dahliae* pathogenicity on tomato. **(a)** Phylogenetic tree of *V. dahliae*. Previously characterized tomato-pathogenic and non-pathogenic strains are shown in bold black and blue, respectively. Strains that were phenoytyped in this study are shown in red. Phylogenetic relationships were inferred using RealPhy (Bertels et al., 2014). *V. alfalfae* strain ms102 was used to root the tree. **(b)** Typical appearance of tomato upon mock-inoculation or inoculation with *V. dahliae* strains JR2, V991, V117, T9, ST100, Vd39 and V574 at 21 days post inoculation (dpi). **(c)** Box plots display canopy area quantifications of inoculated tomato plants from three biological replicates, with dots representing the canopy area of individual plants, the horizontal line within box denotes the median, the box represents the interquartile range, and whiskers extend to minimum and maximum values. Significant differences are indicated (paired Student’s *t*-test; ***: *P*<□0.001; ns: not significant). **(d)** Fungal recovery expressed as the percentage stem slices (n=15) from which *V. dahliae* grew out on potato dextrose agar from three biological experiments shown as mean ± SD (paired Student’s *t*-test; ***: *P* <□0.001; ns: not significant).

#### Comparative genomics identifies pathogenicity effector candidates

Previously, gapless telomere-to-telomere genome assemblies of the tomato-pathogenic strains JR2 and VdLs17 were generated (Faino et al., 2015). Here, we used the genome sequences of nine additional pathogenic strains (2009-605, DVD3, DVD-S161, DVD-31, DVD-S26, DVD-S29, DVD-S94, V4, and CBS38166) and six non-pathogenic strains (ST100, V991, V117, T9, Vd39 and V574) for comparative genome analysis and identified ~133 kb of sequence that is shared by all pathogenic strains and absent from all non-pathogenic strains, collectively encoding thirty-four genes, five of which encode effectors (Figure 2a; Table S3). While four of these are not expressed during host colonization, *VDAG_JR2_Chr3g13460* is expressed with a peak in expression at 12 days post inoculation (Figure 2b), and tentatively named *Tom1*, for potentially mediating *V. dahliae* pathogenicity on <u>tom</u>ato. *Tom1* encodes a secreted protein of 127 amino acids, with no predicted functional domains (InterPro; Finn et al., 2017). To analyse potential *Tom1* gene diversity, we mined the genomes of 39 sequenced *V. dahliae* strains for *Tom1* gene sequence variation. Intriguingly, only one single nucleotide polymorphism (SNP) was identified in one pathogenic strain, which results in a synonymous substitution.

**Figure 2.**
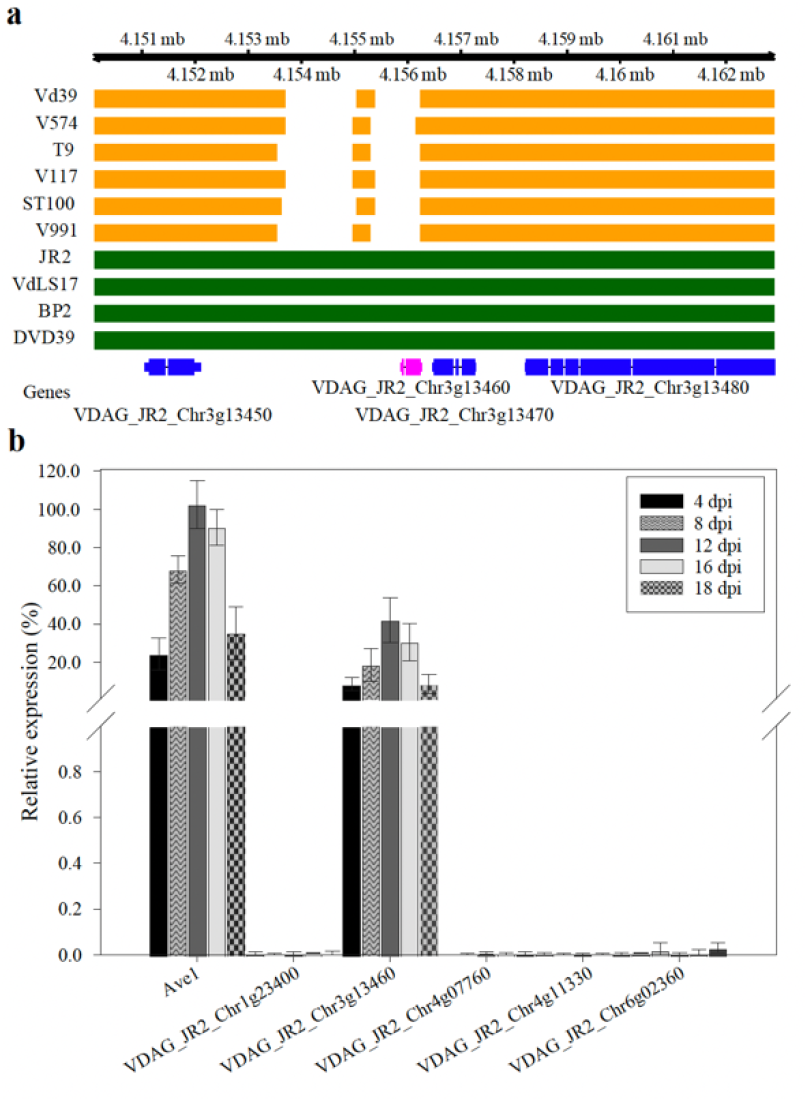
Identification of a putative tomato pathogenicity effector. **(a)** Alignment of contigs of tomato-pathogenic *V. dahliae* strains (green) and strains that do not infect tomato (yellow) showing regions that lack in the non-pathogenic strains. **(b)** Two-week-old tomato seedlings were root-inoculated with *V. dahliae* strain JR2 and were harvested from 4 to 18 days post inoculation (dpi). Real-time PCR was performed to determine the relative expression of five effector genes using *V. dahliae GAPDH* as reference. *Ave1* was used as positive control. Bars represent averages with standard deviation of two biological repeats.

#### The Tom1 effector is essential for *V. dahliae* pathogenicity on tomato

To investigate the role of *Tom1* in tomato pathogenicity, we generated *Tom1* deletion mutants (Figure S1) in *V. dahliae* strain JR2. When compared with wild type JR2, *Tom1* deletion strains (J*Tom1#1* and J*Tom1*#2) lost their ability to infect tomato, as no disease symptoms could be observed (Figure 3a), which was corroborated by canopy area measurements (Figure 3b). Moreover, stem section plating assays failed to recover fungus from stem sections of *ΔTom1-*inoculated plants, whereas the fungus was recovered from JR2-inoculated plants (Figure 3c). Importantly, the loss of pathogenicity displayed by *Tom1* deletion strains was restored in genomic complementation strains (Figure 3a–c). Collectively, these data show that *Tom1* encodes a pathogenicity factor that is crucial for tomato colonization.

**Figure 3.**
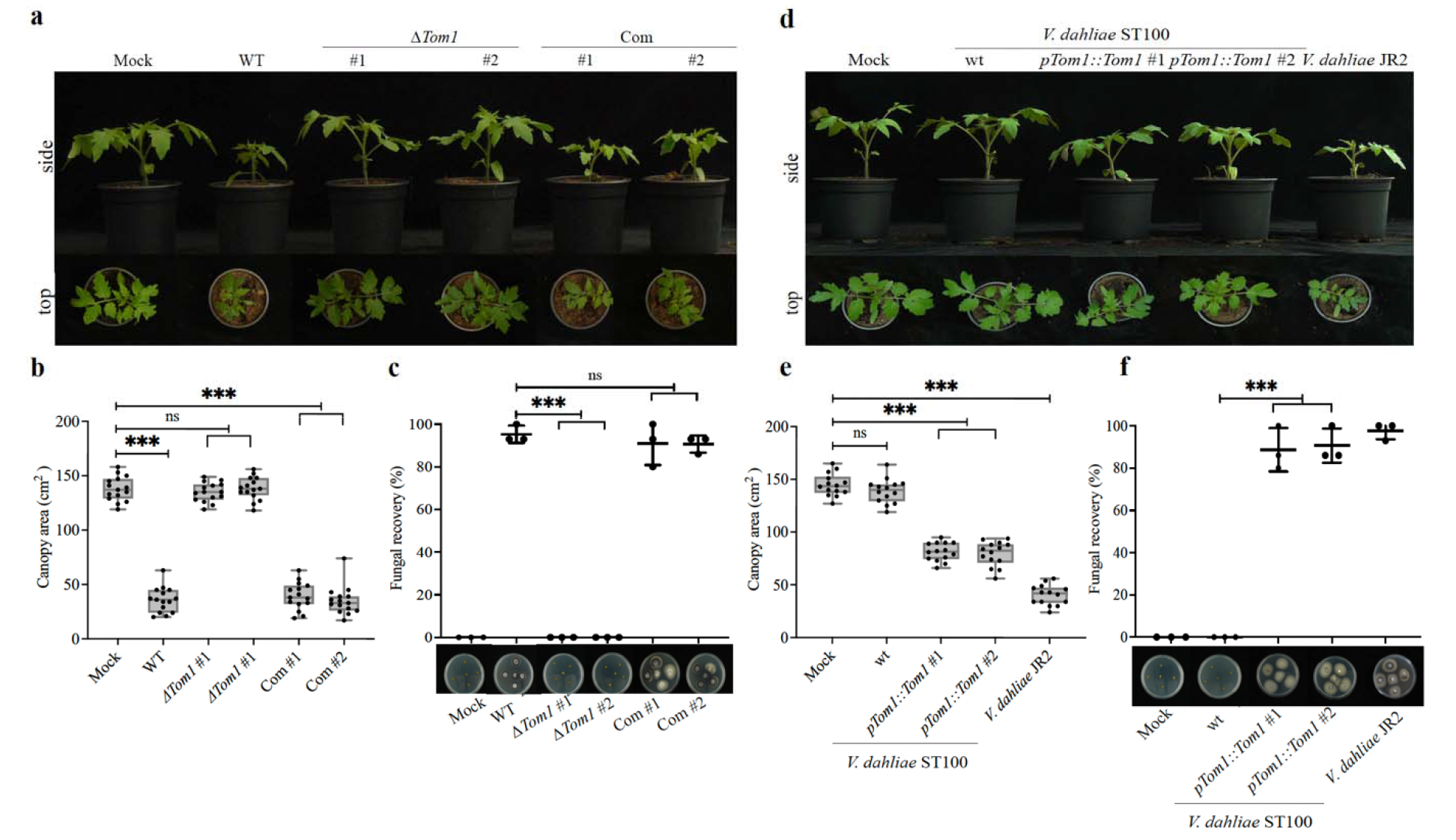
*Tom1* is required and sufficient for *Verticillium dahliae* pathogenicity on tomato. **(a)** Typical appearance of tomato upon mock-inoculation or inoculation with wild type strain JR2 (WT), two independent *Tom1* deletion strains (*ΔTom1#1* and *ΔTom1#2*) and two independent *Tom1* complementation strains (Com#1 and Com#2) at 21 days post inoculation (dpi). (**b**) Box plots display canopy area quantifications of inoculated tomato plants from three biological replicates, with dots representing the canopy area of individual plants, the horizontal line within box denotes the median, the box represents the interquartile range, and whiskers extend to minimum and maximum values. Significant differences are indicated (paired Student’s *t*-test; ***: *P*<□0.001; ns: not significant). **(c)** Fungal recovery expressed as the percentage stem slices (n=15) from which *V. dahliae* grew out on potato dextrose agar from three biological experiments shown as mean ± SD (paired Student’s *t*-test; ***: *P* <□0.001; ns: not significant). **(d)** Typical appearance of tomato upon mock-inoculation or inoculation with wild type strain ST100 (WT), two independent *Tom1* expression transformants (J*Tom1#1* and *ΔTom1#2*) and *V. dahliae* strain JR2 at 21 dpi. **(e)** Box plots display canopy area quantifications of inoculated tomato plants from three biological replicates, with dots representing the canopy area of individual plants, the horizontal line within box denotes the median, the box represents the interquartile range, and whiskers extend to minimum and maximum values. Significant differences are indicated (paired Student’s *t*-test; ***: *P*<□0.001; ns: not significant). **(f)** Fungal recovery expressed as the percentage stem slices (n=15) from which *V. dahliae* grew out on potato dextrose agar from three biological experiments shown as mean **±** SD (paired Student’s *t*-test; ***: *P* <□0.001; ns: not significant).

To investigate whether *Tom1* is not only required, but also sufficient to mediate tomato pathogenicity, we introduced *Tom1* into the non-pathogenic strain ST100 (Figure S1). Interestingly, tomato plants inoculated with these *Tom1* expression strains showed stunting when compared with plants that were inoculated with the wild-type ST100 strain (Figure 4d,e), and fungal outgrowth from plated stem sections could be observed (Figure 4f). Thus, introduction of *Tom1* conferred the ability to cause disease on tomato.

**Figure 4.**
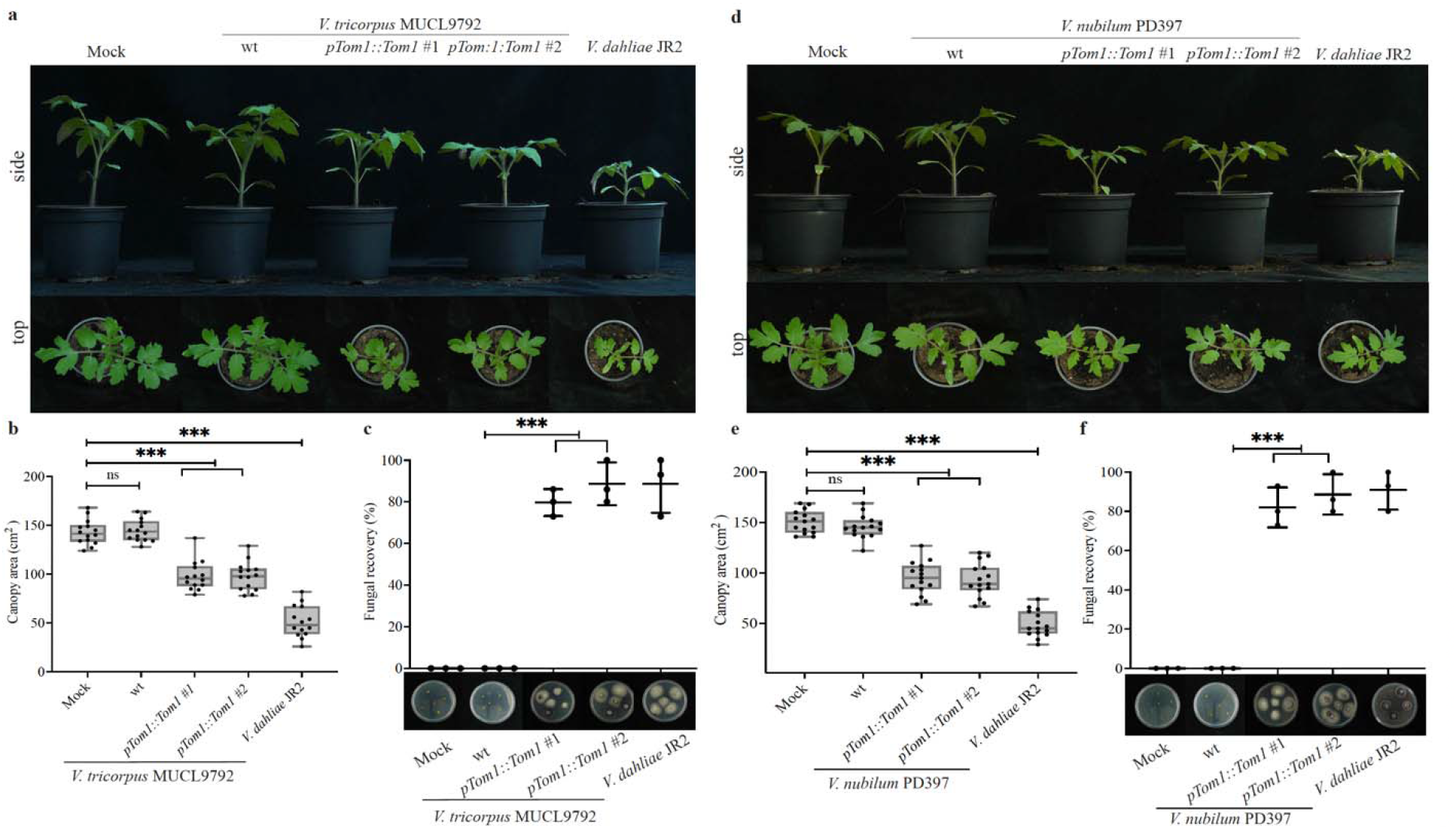
*Tom1* introduction into *Verticillium tricorpus* or *V. nubilum* confers pathogenicity on tomato. a-c: *V. tricorpus* strain MUCL9792, d-f: *V. nubilum* strain PD397 **(a,d)** Typical appearance of tomato upon mock-inoculation or inoculation with wild type *Verticillium* (WT), two independent *Tom1* expression transformants (*pTom1::Tom1#1* and *pToml::Toml#2*) and *V. dahliae* strain JR2 at 21 days post inoculation (dpi). **(b,e)** Box plots display canopy area quantifications of inoculated tomato plants from three biological replicates, with dots representing the canopy area of individual plants, the horizontal line within box denotes the median, the box represents the interquartile range, and whiskers extend to minimum and maximum values. Significant differences are indicated (paired Student’s *t*-test; ***: *P*<□0.001; ns: not significant). **(c,f)** Fungal recovery expressed as the percentage stem slices (n=15) from which *V. dahliae* grew out on potato dextrose agar from three biological experiments shown as mean ± SD (paired Student’s *t*-test; ***: *P* <□0.001; ns: not significant).

#### *Tom1* introduction confers pathogenicity to non-pathogenic sister species

Previous studies have shown that *V. tricorpus* and *V. nubilum* have a saprophytic rather than a pathogenic lifestyle (Inderbitzin et al., 2011; Seidl et al., 2015). Thus, we tested whether *V. tricorpus* strain MUCL9792 and *V. nubilum* strain PD397 can gain tomato pathogenicity upon *Tom1* introduction (Figure S1). As anticipated, wild type *V. tricorpus* and *V. nubilum* were not able to cause disease on tomato (Figure 4a,d). Remarkably, *Tom1-*expressing transformants of both species resulted in pathogenicity on tomato, as evidenced by significant reductions in canopy area development (Figure 4b,e), as well as by fungal recovery assays (Figure 4c,f). Taken together, our data show that the *Tom1* effector is necessary and sufficient to cause tomato disease.

#### Proteomics uncovers *in planta* targets of nuclear-targeted Tom1

The observation that Tom1 contributes to virulence on *Nicotiana benthamiana* (Li, 2019; Li *et al*., unpublished) strongly suggests conservation of molecular virulence targets in this species. Thus, we transiently expressed Tom1 as fusion protein with green fluorescent protein (GFP) in *N. benthamiana* and performed immunopurification followed by mass spectrometry-based identification, using a sequence-unrelated *V. dahliae* effector (Sue1) as control. Principal component analysis (PCA) revealed clear differences in protein composition between immunopurified proteins from leaves expressing the Tom1-GFP fusion protein, the Sue1-GFP fusion protein, or no heterologous protein (Figure 5a). In total, ten proteins were identified that were significantly enriched in samples from leaves expressing GFP-tagged Tom1 that we designated putative interactors PI-1 to PI-10 (Figure 5b, Table S4). These include nuclear transcription factors that belong to the auxin response factor (ARF) family (PI-1 to PI-6) and cytoplasmic pyruvate kinases (PI-7 to PI-10).

**Figure 5.**
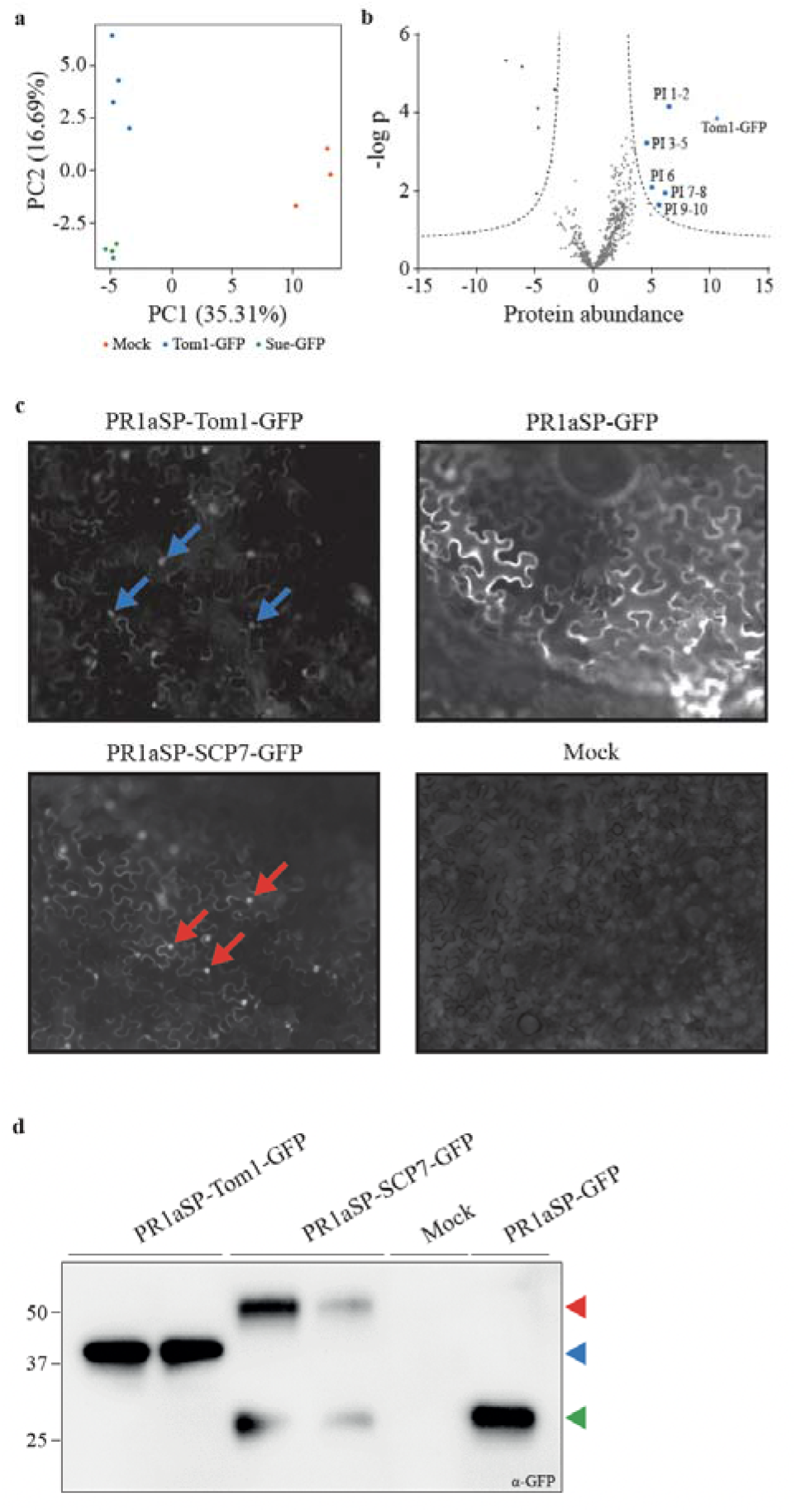
Identification of *in planta* molecular targets of the nuclear-localized Tom1 effector. **(a)** Principal component analysis (PCA) of protein samples containing Tom1-GFP, a sequence-unrelated *V. dahliae* effector (Sue1-GFP) or no heterologously expressed protein (mock) and co-immunopurified plant proteins. **(b)** Volcano plot depicting differential abundance of *N. benthamiana* proteins co-immunopurified with Tom1-GFP or with Sue1-GFP. Blue data points are significantly enriched in Tom1-GFP samples and marked as <u>p</u>utative <u>i</u>nteractor (PI-1 to PI-10) (Student’s T-test, two-tailed; FDR=0.01 and S0=2). The triangle represents Tom1-GFP. **(c)** Live-cell imaging of Tom1-GFP, the nuclear-localiz**ed** *V. dahliae* effector SCP7-GFP, and free GFP in *N. benthamiana* leaf pavement cells at 2 days af**ter** agroinfiltration with a laser-scanning confocal microscope (single optical section). Blue and red arro**ws** indicate nuclei expressing Tom1-GFP and SCP7-GFP, respectively. **(d)** Immunodetection of transiently**-** expressed Tom1-GFP, SCP7-GFP and GFP in *N. benthamiana* with anti-GFP antibody. Expected molecular weights of SCP7-GFP, Tom1-GFP and GFP are indicated by the red, blue and green arrowheads, respectively.

To determine whether the nuclear ARF transcription factors or the cytoplasmic pyruvate kinases are the most likely Tom1 targets, we investigated the subcellular localization of Tom1 *in planta* by expressing Tom1-GFP protein in *N. benthamiana* leaves and performed confocal microscopy. To mimic the situation during *V. dahliae* infection, the protein was expressed with the signal peptide of the *Nicotiana tabacum PR1a* gene for extracellular delivery. Interestingly, in leaves expressing Tom1-GFP, fluorescence was markedly observed in the nucleus of pavement cells, while fluorescence was predominantly detected at their periphery in leaves that expressed free GFP (Figure 5c). This observation suggests that Tom1 is targeted to nucleus. In order to support this finding, we transiently expressed a GFP-tagged version of the *V. dahliae* effector protein SCP7, that was previously shown to localize to host nuclei (Zhang *et al*., 2017; Figure 5c). Extraction of total proteins from the leaves that were subjected to live-cell imaging, followed by immunodetection with anti-GFP antibodies revealed signals matching the expected molecular weights of the expressed proteins (Figure 5d). Collectively, these results indicate that Tom1 is targeted to host nuclei. Therefore, the ARF transcription factors are the most likely *in planta* target proteins of the Tom1 effector, which was further investigated in tomato.

#### Tom1 interacts with tomato ARFs

ARF transcription factors are conserved components of auxin signalling regulate auxin-mediated gene expression (Roosjen *et al*., 2018). ARFs are typically encoded by a gene family, with 23 members in the *Arabidopsis thaliana* genome (Roosjen *et al*., 2018). Phylogenetic analysis of ARF proteins from *N. benthamiana*, tomato, and *A. thaliana* revealed a single ortholog of the NbARF1 protein and two orthologs of the NbARF2 proteins in tomato, which we refer to as SlARF1, SlARF2a and SlARF2b, respectively (Figure S2). Whereas expression of *SlARF1* and *SlARF2a* was detected throughout *V. dahliae* infection of tomato, no expression of *SlARF2b* could be observed (Figure 6a). To assess whether *SlARF1* and *SlARF2a* interact with Tom1 *in planta*, we transiently co-expressed Myc-tagged Tom1 and GFP-tagged SlARF1 or SlARF2a in *N. benthamiana*, followed by immunopurification. Interestingly, Tom1 co-immunopurified with both SlARF proteins, but not with free GFP (Figure 6b), demonstrating that these proteins a**re** targeted by the Tom1 effector *in planta*.

**Figure 6.**
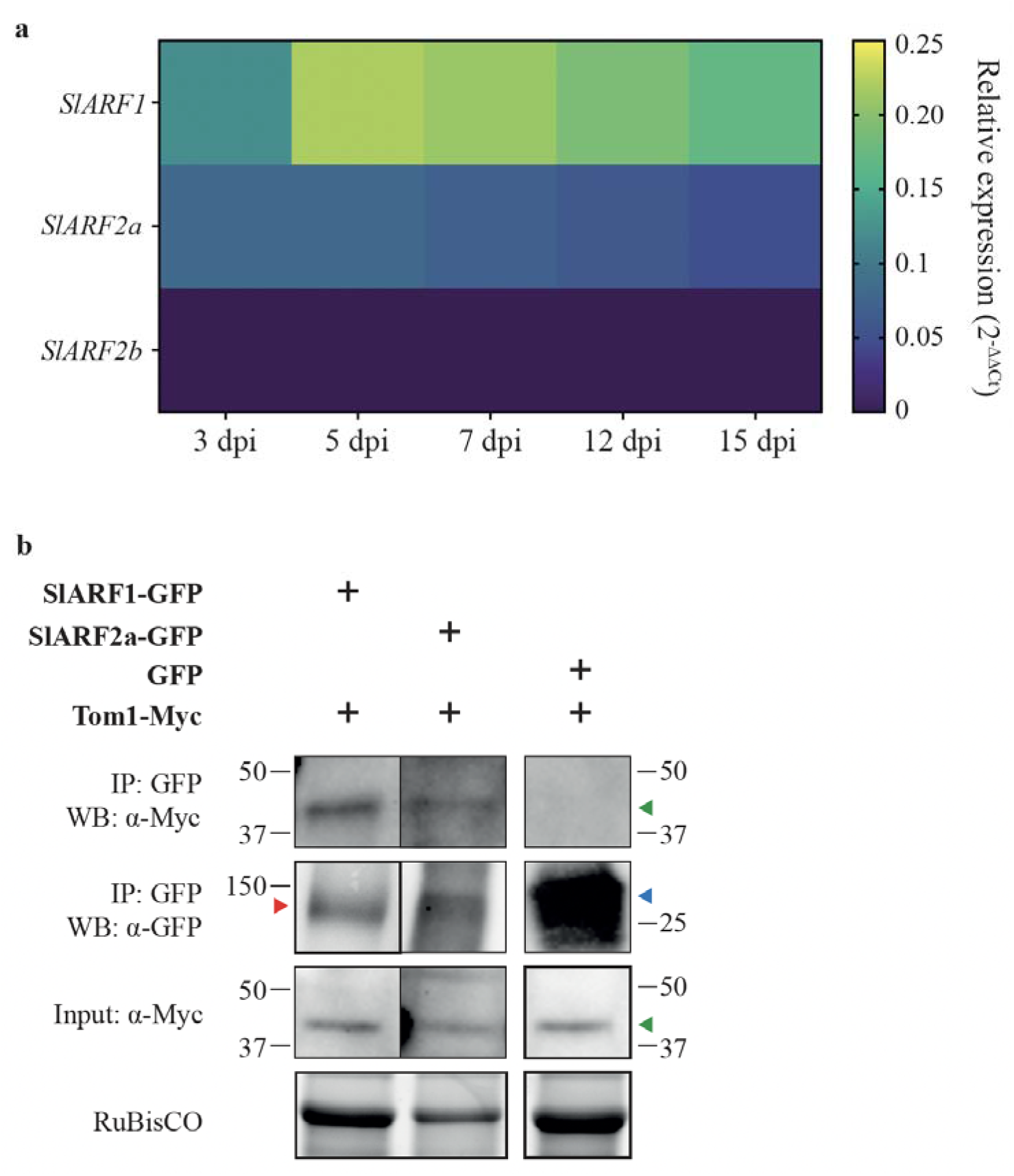
Tomato auxin response factors SlARF1 and SlARF2a targeted by the Tom1 effector. **(a)** Expression of *SlARF1, SlARF2a* and *SlARF2b*, normalized to tomato tubulin (*SlTub*) during *V. dahliae* colonization of tomato from 3 to 15 days post inoculation (dpi). **(b)** Co-immunoprecipitation assays with Myc-tagged Tom1 and GFP-tagged SlARF1 or SlARF2a or free GFP protein as control in *N. benthamiana*. Arrowheads indicate Tom1-Myc (green), SlARF1-GFP and SlARF2a-GFP (red), and GFP (blue).

#### *SlARF2a* silencing impairs tomato colonization

The observations that *SlARF1* and *SlARF2a* are induced during *V. dahliae* colonization, and that Tom1 interacts with these tomato proteins, strongly suggests that SlARF1 and SlARF2a a**re** genuine *in planta* targets of Tom1. To test this, we performed pathogenicity assays on tomato upon repression of *SlARF* expression by means of virus-induced gene silencing (VIGS), using silencing of the β-glucuronidase (*GUS*) gene and the phytoene desaturase (*PDS*) gene as controls. Reduction of *SlARF1* and *SlARF2a* transcript levels was monitored by real time PCR (Figure 7a), whereas *PDS*-silenced plants showed the expected bleaching phenotype (Figure 7b). Subsequently, plants were challenged with *V. dahliae* strain JR2 (WT), a corresponding *Tom1* deletion strain (Δ *Tom1*), or mock-inoculated, and assessed for symptom development and fung**al** colonization. As expected, the *Tom1* deletion strain did not infect any of the inoculated plants (Figure 7c). However, while *SlARF1-silenced* plants inoculated with *V. dahliae* strain JR2 were as stunted as inoculated control plants, *SlARF2a*-silenced plants only showed mild stunting (Figure 7c). Accordingly, whilst *SlARF1-silenced* plants accumulated similar amounts of fungal biomass as control plants, fungal colonization was severely compromised in *SlARF2a*-silenced plants (Figure 7d). Taken together, while *SlARF1* silencing does not affect *V. dahliae* infection, *SlARF2a* silencing drastically impairs tomato colonization.

**Figure 7.**
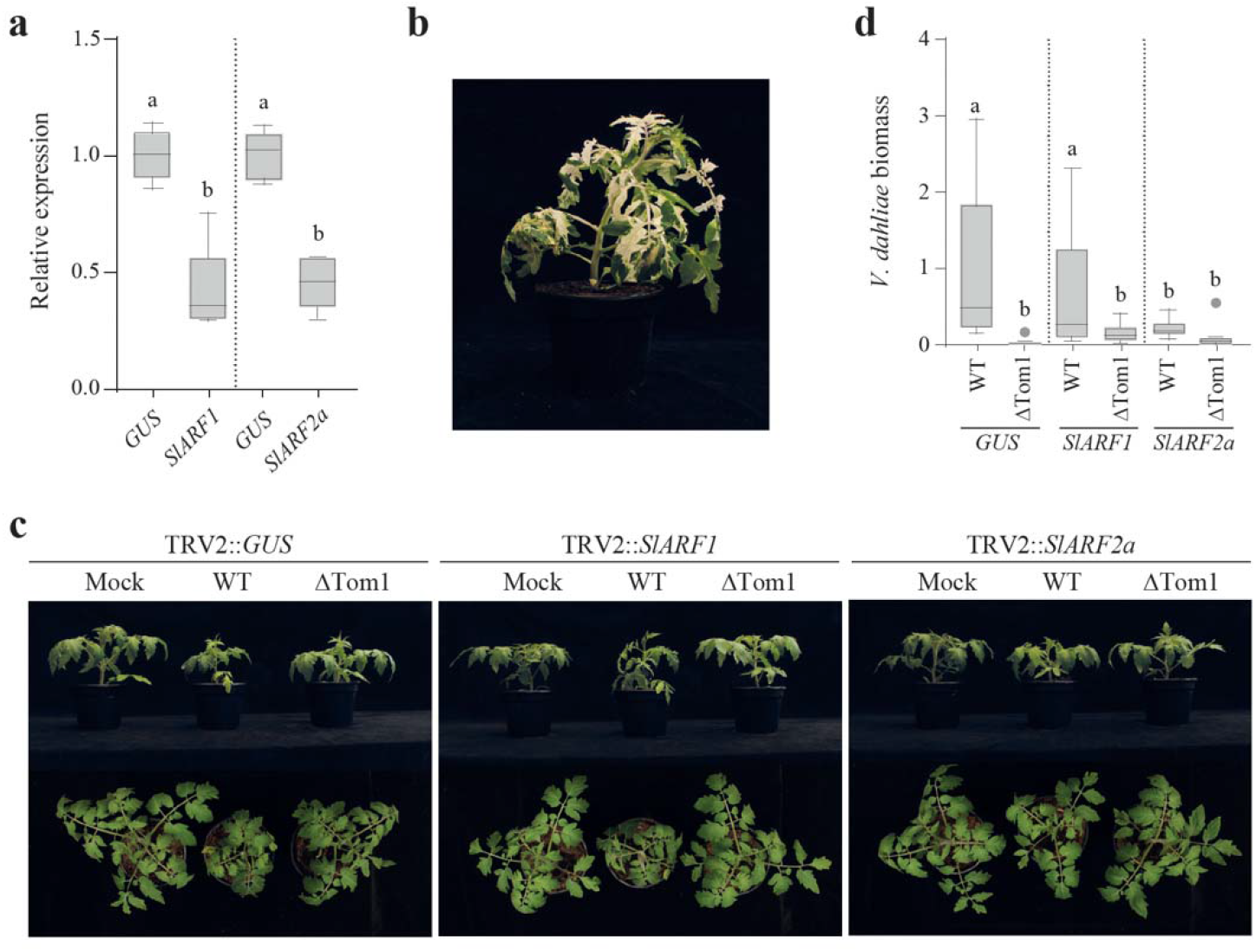
Silencing of *SlARF2a* impairs *Verticillium dahliae* colonization of tomato. **(a)** Transcript levels of *SlARF* genes in *SlARF-* and *GUS*-silenced plants as determined by real time PCR and normalized to tomato Tubulin (*SlTub*). Different letter labels indicate significant differences (one-way ANOVA and Tukey’s multiple comparisons test; n=5 and p<0.05). **(b)** Phenotype of *PDS*-silenced tomato at five weeks after treatment. **(c)** Symptom development in *GUS-* and *SlARF*-silenced tomato plants at three weeks after inoculation with *V. dahliae* strain JR2 or the corresponding *Tom1* deletion strain (*ΔTom?)*. **(d)** *V. dahliae* biomass in *GUS-* and *SlARF*-silenced tomato plants at three weeks after inoculation with *V. dahliae* strain JR2 (WT) or *ΔTom1* determined with real-time PCR by quantification of *V. dahliae* ITS normalized to *SlTub*. Different letter labels indicate significant differences (one-way ANOVA and Tukey’s multiple comparisons test; n=10 and p<0.05).

### DISCUSSION

Plant pathogens typically employ dozens to hundreds of effectors to modulate host immunity and promote disease (Rovenich et al., 2014, Toruño et al., 2016; Wilson and McDowell, 2022). Accordingly, effectors of plant pathogens typically display functional redundancy although exceptions exist where a single effector governs pathogenicity and enables disease establishment (de Jonge et al., 2011; Tan et al., 2015; Win et al., 2012). For example, the transcriptional activator-like effector protein PthA mediates pathogenicity of the bacterial pathogen *Xanthomonas citri* on citrus (Swarup et al., 1991, 1992; Duan et al., 1999), while the ToxA effector of the fungal wheat pathogens *Pyrenophora tritici-repentis* and *Parastagonospora nodorum* confers pathogenicity on wheat lines harbouring *Tsn1* (Friesen et al., 2006). Here, we demonstrate that the Tom1 effector is a pathogenicity factor of *V. dahliae* on tomato, as *Tom1* deletion leads to loss of the ability to colonize tomato plants, whereas its introduction into a non-pathogenic *V. dahliae* isolate, or even into non-pathogenic *Verticillium* sister species, is sufficient to confer the ability to cause disease on tomato.

Understanding the activities of effector proteins is paramount for comprehension of virulence strategies utilized by plant pathogens and, accordingly, effectors can be utilized as probes to identify Achilles’ heels in host plant physiology. Here, we showed that Tom1 interacts with members of the auxin response factor (ARF) family of transcription factors. The observation that the Tom1 effector localizes to the nucleus strongly supports the hypothesis that these transcription factors are indeed targeted by Tom1, which could further be confirmed through VIGS analyses that support the notion that SlARF2a acts as the *V. dahliae* susceptibility factor that is targeted by Tom1. Auxin (indole-3-acetic acid [IAA]) plays major roles in plant growth and development, primarily through regulation of gene transcription, and ARFs are transcription factors that bind as dimers to so-called auxin response elements (AuxREs) in the promoters of auxin-responsive genes (Chapman & Estelle, 2009; Weijers & Wagner, 2016; Leyser, 2018; Roosjen *et al*., 2018). So-called Auxin/Indole-3-Acetic-Acid (Aux/IAA) proteins interact with, and inhibit, the activity of auxin response factors by recruitment of corepressors (Leyser, 2018). In the presence of auxins, Aux/IAA proteins interact with F-box proteins of the Transport Inhibitor Resistant1/Auxin Signalling F-box (TIR1/AFB) family, leading to Aux/IAA ubiquitination and proteasome-mediated degradation, relieving the inhibition of ARFs with subsequent effects on gene expression (Leyser, 2018). Interestingly, auxins have previously also been implicated in plant disease and resistance (Robert-Seilaniantz *et al*., 2011; Kazan & Lyons, 2014; Han & Kahmann, 2019). Whereas auxin signalling promotes resistance against the necrotrophic pathogens *Plectosphaerella cucumerina* and *Botrytis cinerea* in *Arabidopsis thaliana* (Llorente *et al*., 2008), *Pseudomonas syringae* bacteria produce auxin to promote disease development (McClerklin *et al*., 2018; Navarro *et al*., 2006). Similarly, auxin application on rice (*Oryza sativa*) aggravates infections of *Magnaporthe oryzae* and *Xanthomonas oryzae* pv oryzicola (*Xoc*) (Fu *et al*., 2011), suggesting that auxin promotes infections of (hemi)biotrophic pathogens. Auxins have also been implicated in susceptibility to vascular pathogens. While auxin receptor and transporter mutants of *Arabidopsis* show enhanced resistance to *V. dahliae* (Fousia *et al*., 2018), and deletion of the vacuolar auxin transporter WAT1 resulted in enhanced *V. dahliae* resistance in tomato (Hanika, 2020), auxin signalling and transport mutants showed enhanced resistance against *Fusarium oxysporum* in *Arabidopsis* (Kidd *et al*., 2011; Lyons *et al*., 2015).

The relevance of auxin for plant-pathogen interactions is further substantiated by the identification of pathogen effectors that target auxin signalling (Kazan & Lyons, 2014; Kunkel & Johnson, 2021). For example, while *P. syringae* exploits the AvrRpt2 effector protein to promote degradation of Aux/IAA proteins to activate auxin responses (Chen *et al*., 2007; Cui *et al*., 2013), *Phytophthora parasitica* secretes the RxLR effector PSE1 to affect the distribution of auxin transporters to alter local auxin levels in roots (Evangelisti *et al*., 2013). Furthermore, the *Ustilago maydis* Nkd1 effector promotes auxin signalling in maize by targeting transcriptional co-repressors of the Topless/Topless-related (TPL/TPRs) family, resulting in impairment of PAMP-triggered immunity (PTI) (Navarrete *et al*., 2022). To our knowledge, *V. dahliae* Tom1 is the only fungal effector that directly targets ARF transcription factors identified to date.

Considering that ARF1 and ARF2 exhibit only partial functional redundancy in *A. thaliana* (Li *et al*., 2004; Okushima *et al*., 2005; Ellis *et al*., 2005), the differential impact of *SlARF1* and *SlARF2a* gene silencing on *V. dahliae* biomass accumulation in tomato may indicate that *V. dahliae* colonization is favoured by targeting processes regulated by SlARF2a rather than by SlARF1. However, alternatively, SlARF2a may be able to compensate for *SlARF1* silencing whereas the reverse may not occur, as previously observed for Arabidopsis ARF7 and ARF19 (Okushima *et al*., 2005). Finally, we also acknowledge that the residual SlARF1 that remains after silencing might still be sufficient to support Tom1-mediated host colonization, and only genuine *SlARF1* deletion mutants will clarify whether SlARF1 is a target of Tom1 to support *V. dahliae* colonization.

Although auxin affects plant-pathogen interactions through a diversity of mechanisms (Robert-Seilaniantz *et al*., 2011; Kazan & Lyons, 2014; Ma & Ma, 2016; Kunkel & Johnson, 2021), how Tom1-mediated SlARF2a manipulation mediates *V. dahliae* colonization of tomato remains elusive. As *V. dahliae* is not known to promote morphogenesis, such as for accommodation of sheltering and feeding structures by *Agrobacterium tumefaciens* or *Meloidogyne incognita* (Grunewald *et al*., 2009; Kyndt *et al*., 2016), manipulation of auxin signalling to promote abnormal plant cell division is not expected. However, auxins also affect plant immunity through crosstalk with signalling networks of defence-related hormones (Robert-Seilaniantz et al., 2011). Furthermore, given that Arabidopsis *arf2* mutants do not differentially express particular auxin-regulated genes, yet show pleiotropic phenotypes, ARF2 may bind promoters of genes that are not directly regulated by auxin, or interact with proteins that are not directly involved in auxin signalling (Okushima *et al*., 2005; Ellis *et al*., 2005; Lim *et al*., 2010). Future identification of SlARF2a target genes and of SlARF2a-interacting proteins, in presence and absence of the Tom1 effector, may reveal how SlARF2a modulation affects tomato susceptibility to *V. dahliae*.

Thus far, the most sustainable strategy to combat Verticillium wilt disease is the deployment of resistant cultivars (Fradin and Thomma, 2006). However, the two currently deployed resistance genes *Ve1* and *V2* have rapidly been overcome by resistance-breaking strains that continue to threat tomato production (Schaible et al., 1951; Kawchuk et al., 2001; Pegg and Brady, 2002; Usami *et al*, 2017; Chavarro-Carrero et al., 2021), and attempts to discover additional major resistance genes in tomato against *V. dahliae* have failed (Vermeulen et al., 2022). Considering that the impairment of host genes that are exploited by pathogens for disease establishment, so-called disease susceptibility (*S*) genes, is an alternative approach to obtain pathogen resistance (Pavan et al., 2010; van Schie and Takken, 2014), targeting of *SlARF2* may be considered for disease management. Furthermore, pathogen effectors can be used to probe plant germplasm for recognition specificities that can be deployed in resistance breeding (Laugé et al., 1998; Vleeshouwers and Oliver, 2014). Accordingly, Tom1 may be used to screen tomato germplasm for recognition specificities that, if found, may be difficult to overcome for *V. dahliae*, given that Tom1 is a pathogenicity factor on tomato, and that *V. dahliae* overcame *Ve1* and *V2* resistance only through purging the corresponding effector genes (de Jonge et al., 2012; Chavarro-Carrero et al., 2021). Thus, tomato resistance that is based on *Tom1* recognition may be more durable than the resistances that have been deployed thus far.

## ACKNOWLEDGEMENTS

JLL and GLF acknowledge PhD fellowships from the China Scholarship Council (CSC) and the Coordination for the Improvement of Higher Education Personnel (CAPES) from the federal government of Brazil, respectively. Bert Essenstam (Unifarm) is thanked for excellent plant care. BPHJT acknowledges funding by the Alexander von Humboldt Foundation in the framework of an Alexander von Humboldt Professorship endowed by the German Federal Ministry of Education and is furthermore supported by the Deutsche Forschungsgemeinschaft (DFG, German Research Foundation) under Germany’s Excellence Strategy - EXC 2048/1 - Project ID: 390686111 and the Research Council for Earth and Life Sciences (ALW) of the Netherlands Organization for Scientific Research (NWO).

## AUTHOR CONTRIBUTIONS

LF and BPHJT conceived the project. LF, JL, GLF, MFS and BPHJT designed the experiments, LF, JL, GLF, SB, AS and GCMB and performed the experiments and analyzed the data. JL, GLF, and BPHJT wrote the manuscript. All authors read and approved the final manuscript.

## DATA AVAILABLILITY

Data sharing is not applicable to this article as no new data were created or analyzed in this study.

