## Supplemental Information for "A single *Verticillium dahliae* effector determines pathogenicity on tomato by targeting auxin response factors"

### SUPPLEMENTARY INFORMATION

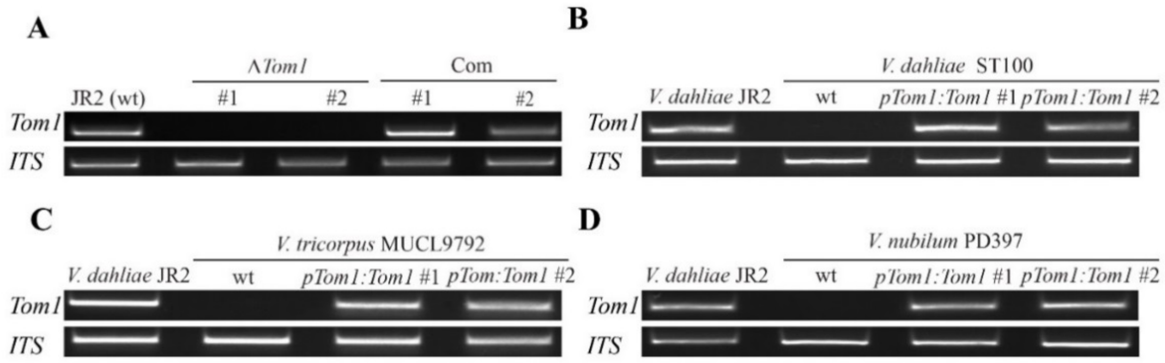


**Figure S1. Verification of *Tom1* deletion and complementation strains. (A)** Amplification of *Tom1* from genomic DNA of *V. dahliae* strain JR2, two independent *Tom1* deletion strains (*ΔTom1*#1 and *ΔTom1*#2) and two *Tom1* complementation strains (Com#1 and Com#2). As an endogenous control, a fragment of the *Verticillium* ITS region was amplified. **(B)** Amplification of *Tom1* from genomic DNA of *V. dahliae* strain JR2, wild type *V. dahliae* strain ST100 (wt), and two independent *Tom1* expression transformants (*pTom1::Tom1*#1 and *pTom1::Tom1*#2). As an endogenous control, a fragment of the *Verticillium* ITS region was amplified. **(C)** Amplification of *Tom1* from genomic DNA of *V. dahliae* strain JR2, wild type *V. tricorpus* strain MUCL9792 (wt), and two independent *Tom*1 expression transformants (*pTom1::Tom1*#1 and *pTom1::Tom1*#2). As an endogenous control, a fragment of the *Verticillium* ITS region was amplified. **(D)** Amplification of *Tom1* from genomic DNA in *V. dahliae* strain JR2, wild type *V. nubilum* strain PD397 (wt), and two independent *Tom1* expression transformants (*pTom1::Tom1*#1 and *pTom1::Tom1*#2). As an endogenous control, a fragment of the *Verticillium* ITS region was amplified.

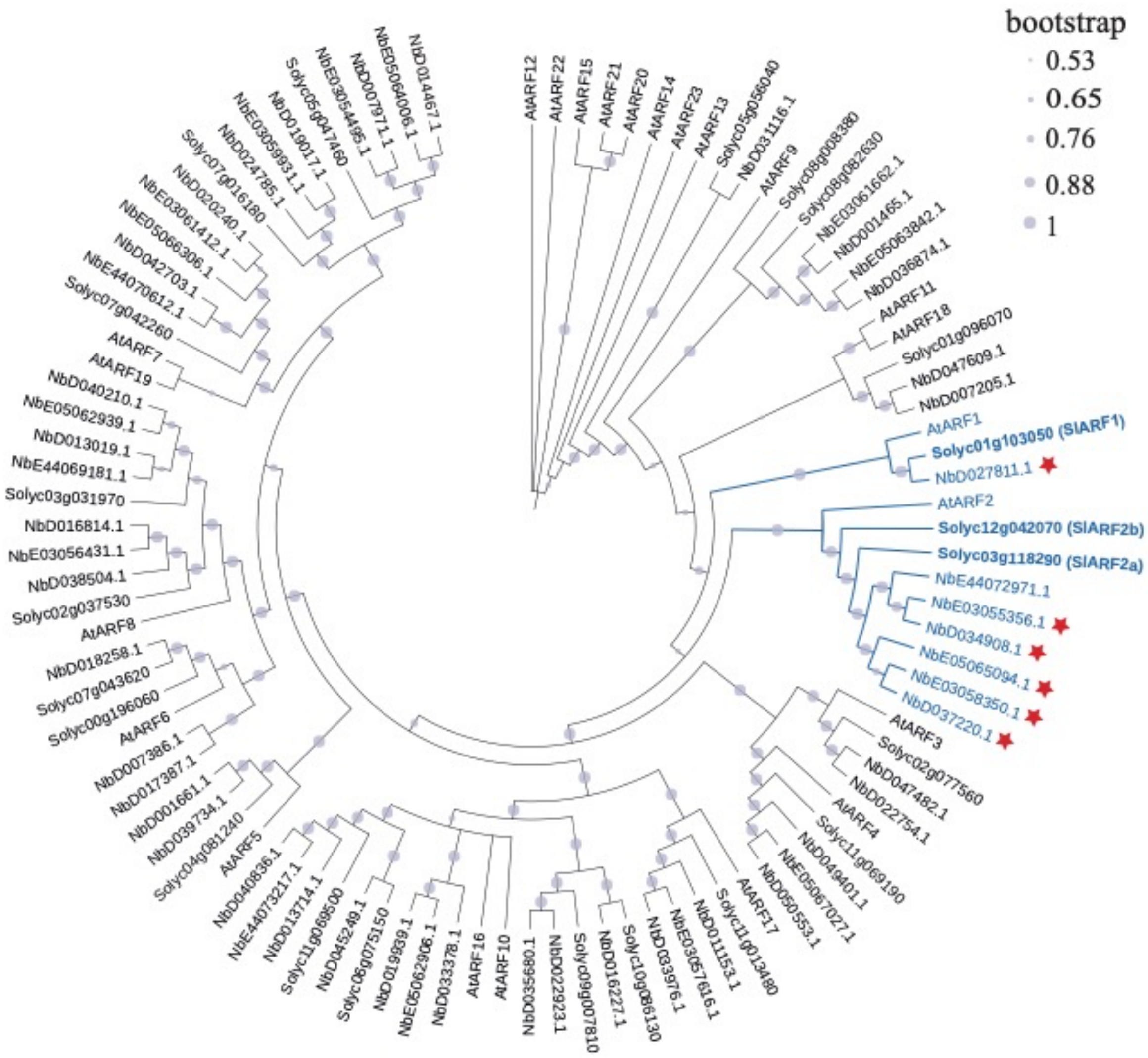


**Figure S2. Phylogeny of ARF transcription factors of Arabidopsis, tomato and *Nicotiana benthamiana*.** *N. benthamiana* proteins identified as putative interactors of the Tom1 effector in the proteomics data (PI1-6) are marked with a red star, and clustering shows that SlARF1, SlARF2a and SlARF2b, which are highlighted in bold, are their most likely tomato orthologs.

**Table S1. Primers used in this study.**

| Name | Oligonucleotide Sequence (5'-3') | Description |
|---|---|---|
| Tom1-LBF | ggtcttaauTGTAATCGGCGATAGGAAGG | *Tom1* deletion, left border |
| Tom1-LBR | ggcattaauGACTTTGGCAACATGGTGAA | *Tom1* deletion, left border |
| Tom1-RBF | ggacttaauAGATCACCCACCTGACCTTG | *Tom1* deletion, right border |
| Tom1-RBR | gggtttaauCGCGACTCTGCCTTCTTAAC | *Tom1* deletion, right border |
| Tom1-com-F | ggggacagctttcttgtacaaagtggaaTGTAATCGGCGATAGGAAGG | *Tom1* complementation |
| Tom1-com-R | ggggacaactttgtataataaagttgtCGCGACTCTGCCTTCTTAAC | *Tom1* complementation |
| Sue1_pDONR_F | ggggacaagtttgtacaaaaaagcaggcttgATTGCCATCCCGCAATCCG | Cloning Sue1 effector |
| Sue1_pDONR_R | ggggaccactttgtacaagaaagctgggtcAAGCTGGCCTGCGTCAAAAG | Cloning Sue1 effector |
| TOM_pDONR_F | ggggacaagtttgtacaaaaaagcaggcttgCTTTCTATCGATAGTCGTCAAGAC | Cloning Tom1 effector |
| TOM_pDONR_R | ggggaccactttgtacaagaaagctgggtcGTGCTCAAGCCTCACTCC | Cloning Tom1 effector |
| PR1a_SP_pDONR_F | ggggacaagtttgtacaaaaaagcaggcttgATGGGATTTGTTCTCTTTTCACAAT | Amplification of PR1a signal peptide |
| PR1a_SP_pDONR_Sue1_R | cggtatcggattgcgggatggcaatGGCACGGCAAGAGTGGG | Fusion PR1a signal peptide with Sue1 effector |
| PR1a_SP_pDONR_TOM_R | cgtcttgacgactatcgatagaaagGGCACGGCAAGAGTGGG | Fusion PR1a signal peptide with Tom1 effector |
| Solyc01g103050_F | ggggacaagtttgtacaaaaaagcaggcttgATGGCACACGTTGCTGC | Cloning SlARF1 |
| Solyc01g103050_R | ggggaccactttgtacaagaaagctgggtcTGCTTTTTCATTGTTGCCAAC | Cloning SlARF1 |
| Solyc03g118290_F | ggggacaagtttgtacaaaaaagcaggcttgATGGCTGCTTCGGAGGTG | Cloning SlARF2a |
| Solyc03g118290_R | ggggaccactttgtacaagaaagctgggtcAGATTCTGCTTGACCAGATTCAG | Cloning SlARF2a |
| Solyc01g103050_VIGS_F | ggggacaagtttgtacaaaaaagcaggcttgATGGCACACGTTGCTGC | Cloning SlARF1 |
| Solyc01g103050_VIGS_R | ggggaccactttgtacaagaaagctgggtcAGTTATCTGTGCATACACCTCATC | Cloning SlARF1 |
| Solyc03g118290_VIGS_F | ggggacaagtttgtacaaaaaagcaggcttgATGGCTGCTTCGGAGGTG | Cloning SlARF2a |
| Solyc03g118290_VIGS_R | ggggaccactttgtacaagaaagctgggtcATTCACAACACGGCATAGAATC | Cloning SlARF2a |
| VdAve1-F (RT) | AGCTTTCTACGCTTGGA | Real-time PCR |
| VdAve1-R (RT) | TTGGCTGGGATTGCT | Real-time PCR |
| VDAG_JR2_Chr1g23400-F | ACTGGAAGGTGACGATGTCC | Real-time PCR |
| VDAG_JR2_Chr1g23400-R | GGGGTACGCTACCTTCTTCC | Real-time PCR |
| VDAG_JR2_Chr3g13460-F | GCTTCACTGTGTCCGCTGTA | Real-time PCR |
| VDAG_JR2_Chr3g13460-R | TCCATATCTTCCCAGGTTGG | Real-time PCR |
| VDAG_JR2_Chr4g07760-F | ACCATGTAAGACTTGAGGCCA | Real-time PCR |
| VDAG_JR2_Chr4g07760-R | AGCGCTCCTGACTGTTATGA | Real-time PCR |

| Name | Oligonucleotide Sequence (5'-3') | Description |
|---|---|---|
| VDAG_JR2_Chr4g11330-F | GCGTACCTCGATTCTCTTCG | Real-time PCR |
| VDAG_JR2_Chr4g11330-R | TTCCGTAAGGTGGCTGAGAT | Real-time PCR |
| VDAG_JR2_Chr6g02360-F | TTAACATGACGGACCACGAA | Real-time PCR |
| VDAG_JR2_Chr6g02360-R | ACCCTGAAGCACAGCTTCTC | Real-time PCR |
| SlARF1_qPCR_F | TCATGACTGGGACCCTCTTC | Real-time PCR |
| SlARF1_qPCR_R | GACGGGTTATCACCAACACC | Real-time PCR |
| SlARF2a_qPCR_F | ACAAGGAGCTGCACAAAGGT | Real-time PCR |
| SlARF2a_qPCR_R | CCAACAAGCATCATGTCACC | Real-time PCR |
| SlARF2b_qPCR_F | CCCCGATTCACTCAACAAGT | Real-time PCR |
| SlARF2b_qPCR_R | CCAGGATGAGAGGTTTGGAA | Real-time PCR |
| ITS1-F | AAAGTTTTAATGGTTCGCTAAGA | *V. dahliae* biomass quantification |
| ITS1-R | CTTGGTCATTTAGAGGAAGTAA | *V. dahliae* biomass quantification |
| SIRub_F | GAACAGTTTCTCACTGTTGAC | *S. lycopersicum* RuBisCO, real-time PCR |
| SIRub_R | CGTGAGAACCATAAGTCACC | *S. lycopersicum* RuBisCO, real-time PCR |
| SlTub_F | AACCTCCATTCAGGAGATGTTT | *S. lycopersicum* Tubulin, real-time PCR |
| SlTub_R | TCTGCTGTAGCATCCTGGTATT | *S. lycopersicum* Tubulin, real-time PCR |

**Table S2.** ***Verticillium dahliae*** **strains used in this study.**

| Strain | Platform | Reference | Origin | Location |
|---|---|---|---|---|
| HN | Illumina | Xu et al., 2012 | Cotton | China |
| cd3 | Illumina | Xu et al., 2012 | Cotton | China |
| VdLs17 | PacBio | Faino et al., 2015 | Lettuce | USA |
| JR2 | PacBio | Faino et al., 2015 | Tomato | Canada |
| Vd57 | Illumina | Chavarro-Carrero et al., 2021 | Strawberry | Germany |
| V152 | Illumina | Kombrink et al., 2017 | Oak | Hungary |
| Vd52 | Illumina | Kombrink et al., 2017 | Pepper | Austria |
| van Dijk | Illumina | Kombrink et al., 2017 | Chrysanthemum | The Netherlands |
| BP2 | Illumina | Zhang et al., 2012 | Cotton | China |
| ST16.01 | Illumina | Chavarro-Carrero et al., 2021 | Cotton | Syria |
| 2009-605 | Illumina | Chavarro-Carrero et al., 2021 | Bell pepper | Ukraine |
| V4 | Illumina | Keykhasaber, 2017 | Olive | Spain |
| CBS38166 | Illumina | de Jonge et al., 2012 | Tomato | Canada |
| DVD-S26 | Illumina | de Jonge et al., 2012 | Soil | Canada |
| VdLS16 | Illumina | de Jonge et al., 2012 | Lettuce | USA |
| ST14.01 | Illumina | de Jonge et al., 2012 | Pistachio | USA |
| DVD-S29 | Illumina | de Jonge et al., 2012 | Soil | Canada |
| DVD-31 | Illumina | de Jonge et al., 2012 | Tomato | Canada |
| DVD-161 | Illumina | de Jonge et al., 2012 | Tomato | Canada |
| DVD-S94 | Illumina | de Jonge et al., 2012 | Soil | Canada |
| DVD-3 | Illumina | de Jonge et al., 2012 | Potato | Canada |
| JKG8 | Illumina | Kombrink et al., 2017 | Potato | The Netherlands |
| 85S | PacBio | Chavarro-Carrero et al., 2021 | Sunflower | France |
| Vd39 | Illumina | Chavarro-Carrero et al., 2021 | Sunflower | Germany |
| V574 | Illumina | Milgroom et al., 2014 | Artichoke | Spain |
| v700 | Illumina | Milgroom et al., 2014 | Artichoke | Spain |

| | | | | |
|---|---|---|---|---|
| v679 | Illumina | Milgroom et al., 2014 | Artichoke | Spain |
| T9 | Illumina | Keykhasaber, 2017 | Cotton | USA |
| TM6 | Illumina | Keykhasaber, 2017 | Cotton | China |
| V117 | Illumina | Keykhasaber, 2017 | Olive | Spain |
| V991 | Illumina | Zhang et al., 2012 | Cotton | China |
| CQ2 | PacBio | Chavarro-Carrero et al., 2021 | Cotton | China |
| ST100 | Illumina | de Jonge et al., 2012 | Soil | Belgium |
| 463 | Illumina | Chavarro-Carrero et al., 2021 | Cotton | Mexico |

**Table S3. Tomato-pathogenic strain-specific genes identified by comparative genomics.**

| Gene ID | Effector |
|---|---|
| VDAG_JR2_Chr1g01610 | No |
| VDAG_JR2_Chr1g01620 | No |
| VDAG_JR2_Chr1g07050 | No |
| VDAG_JR2_Chr1g07060 | No |
| VDAG_JR2_Chr1g12270 | No |
| VDAG_JR2_Chr1g13280 | No |
| VDAG_JR2_Chr1g23400 | Yes |
| VDAG_JR2_Chr2g00380 | No |
| VDAG_JR2_Chr2g00780 | No |
| VDAG_JR2_Chr2g03850 | No |
| VDAG_JR2_Chr2g03860 | No |
| VDAG_JR2_Chr2g12810 | No |
| VDAG_JR2_Chr3g06060 | No |
| VDAG_JR2_Chr3g09230 | No |
| VDAG_JR2_Chr3g09240 | No |
| VDAG_JR2_Chr3g09250 | No |
| VDAG_JR2_Chr3g13460 | Yes |
| VDAG_JR2_Chr4g02130 | No |
| VDAG_JR2_Chr4g07760 | Yes |
| VDAG_JR2_Chr4g11330 | Yes |
| VDAG_JR2_Chr5g02370 | No |
| VDAG_JR2_Chr5g08390 | No |
| VDAG_JR2_Chr6g02360 | Yes |
| VDAG_JR2_Chr6g02370 | No |
| VDAG_JR2_Chr6g03370 | No |
| VDAG_JR2_Chr6g03410 | No |
| VDAG_JR2_Chr7g03300 | No |
| VDAG_JR2_Chr7g05290 | No |
| VDAG_JR2_Chr8g06110 | No |
| VDAG_JR2_Chr8g06120 | No |
| VDAG_JR2_Chr8g08950 | No |
| VDAG_JR2_Chr8g09620 | No |
| VDAG_JR2_Chr8g09630 | No |
| VDAG_JR2_Chr8g09640 | No |

**Table S4. List of putative *in planta* targets of the Tom1 effector.**

| Putative Interactor | Protein ID | Predicted annotation | Predicted localization |
|---|---|---|---|
| PI-1 | NbD034908.1 | Auxin response factor 2 | Nucleus |
| PI-2 | NbE03055356.1 | Auxin response factor 2 | Nucleus |
| PI-3 | NbD037220.1 | Auxin response factor 2 | Nucleus |
| PI-4 | NbE05065094.1 | Auxin response factor 2 | Nucleus |
| PI-5 | NbE03058350.1 | Auxin response factor 2 | Nucleus |
| PI-6 | NbD027811.1 | Auxin response factor 1 | Nucleus |
| PI-7 | NbD048621.1 | Pyruvate kinase | Cytoplasm |
| PI-8 | NbD010296.1 | Pyruvate kinase | Cytoplasm |
| PI-9 | NbE03054249.1 | Pyruvate kinase 1 | Cytoplasm |
| PI-10 | NbD025603.1 | Pyruvate kinase 1 | Cytoplasm |

### SUPPLEMENTARY METHODS

#### Gene expression analysis

Two-week-old tomato (cv. Moneymaker) seedlings were inoculated with *V. dahliae* strain JR2 and stems harvested at designated time points. Next, samples were grinded to a fine powder using mortar and pestle. Approximately 100 mg of each sample was used to perform RNA isolation using the Spectrum Plant Total RNA kit (Sigma, Saint Louis, USA) according to the manufacturer's instructions. The resulting RNA samples were quantified spectrophotometrically (NanoDrop, Thermo Fisher Scientific, California, USA) and checked for integrity by gel electrophoresis, after which they were reverse transcribed using the M-MLV reverse transcriptase (Promega, Leiden, the Netherlands) according to the manufacturer's instructions. The resulting cDNAs were used for quantification of transcript levels by means of real-time PCR in a C1000 Touch™ Thermal Cycler (Bio-Rad, California, USA) using a SYBR green master mix kit (Bioline, London, UK) according to the manufacturer's instructions. Gene expression was normalized to *SlTub* (Table 2). $C_t$ values were analyzed with the $2^{(\text{-Delta Delta Ct})}$ method (Livak & Schmittgen, 2001) and plotted using GraphPad Prism (version 9.2.0).

#### Proteomics data analysis

Peptide spectra were searched in MaxQuant (version 1.6.17.0) using the Andromeda search engine (Cox *et al.*, 2011) with default settings and label-free quantification against the NbDE *N. benthamiana* proteome dataset (Kourelis *et al.*, 2019) and including a dataset of frequently occurring contaminants. Protein quantification was performed in MaxQuant (Tyanova *et al.*, 2016a), based on unique and razor peptides including all modifications. Identified protein groups were then subjected to analysis using Perseus (version 1.6.12.0) (Tyanova *et al.*, 2016b). First, reverse and contaminant proteins as well as those only identified by matching were filtered out. Next, only those protein groups identified in 3 out of 4 biological replicates of either Tom1-GFP or Sue1-GFP were selected. The LFQ values were log2 transformed, and missing values were imputed using a normal distribution. A both-sided Student's T test was used and adjusted using a permutation-based adjustment (FDR=0.05, 250 randomizations) to identify protein groups differentially enriched in Tom1-GFP samples.

#### Live-cell imaging by confocal microscopy

For determination of protein subcellular localization *in planta*, leaves of six-week-old *Nicotiana benthamiana* plants were transiently transformed by *Agrobacterium*-mediated expression with the PR1aSP-Tom1::pSOL2095, PR1aSP::pSOL2095 or the PR1aSP-SCP7::pSOL2095 construct (Zhang *et al.*, 2017) and the P19 silencing suppressor in a 1:1 ratio at a final optical density ($OD_{600}$) of 0.8 for each construct. Two days after infiltration, leaf pieces were cut and mounted on glass slides for confocal microscopy using a Leica CLSM SP8 microscope (Leica, Wetzlar, Germany). The GFP was excited at 488 nm. Integrity of GFP-tagged proteins was assessed by western blotting as described previously (Liebrand *et al.*, 2012) using αGFP-HRP antibody (Miltenyi Biotec GmbH, Bergisch Gladbach, Germany).

**Virus-induced gene silencing (VIGS) in tomato seedlings**

VIGS was performed using a tobacco rattle virus (TRV)-based system as described previously (Liu *et al.*, 2002). To generate silencing constructs, exonic 150-300 bp regions of each of the *SlARF* genes were amplified by PCR using genomic DNA obtained from tomato as template. The resulting amplicons were cloned into a Gateway compatible version of the pTRV2 plasmid (Liu *et al.*, 2002) and the clones that were obtained were confirmed by Sanger sequencing. Confirmed clones were transformed by electroporation into *A. tumefaciens* strain GV3101 for transient transformation of ten-day-old tomato seedlings by means of cotyledon infiltration. Silencing of the non-endogenous β-glucuronidase (GUS) gene was carried out as control. Additionally, silencing of the tomato phytoene desaturase (*PDS*) gene was included as control, as efficient silencing of this gene leads to a clear bleaching phenotype. At 14 days after silencing, plants were inoculated with conidiospores of *V. dahliae* strain JR2, a deletion strain lacking the *Tom1* gene (Δ*Tom1*) or water (mock) by means of root dipping as described previously (Fradin *et al.*, 2009). Silencing levels of the genes that were targeted and the relative amount of fungal biomass in inoculated plants were determined at 21 days after inoculation through real-time PCR in a C1000 Touch™ Thermal Cycler (Bio-Rad, California, USA) using a SYBR green master mix kit (Bioline, London, UK) according to the manufacturer's instructions. Gene expression was normalized to *SlTub* and *V. dahliae* biomass was quantified using specific primers targeting the internal transcribed spacer (ITS) region of the *V. dahliae* ribosomal DNA (Table 2). $C_t$ values were analyzed with the $2^{(-\mathrm{Delta\ Delta\ Ct})}$ method (Livak & Schmittgen, 2001) and plotted using GraphPad Prism (version 9.2.0).

**Phylogenetic analysis of ARF transcription factors**

To determine the tomato orthologs of the *Nicotiana benthamiana* ARF proteins that were identified by mass spectrometry, the predicted proteomes of tomato (version 4.0; Solanaceae Genomics Network), *N. benthamiana* (NbDE dataset; Kourelis *et al.*, 2019) and *Arabidopsis thaliana* (version 10; TAIR) were queried for proteins containing the auxin response factor domain (IPR010525) using HMMer (version 3.3.2) (Eddy, 2011) and a local Pfam database (version 35.0) (Mistry *et al.*, 2021). Retrieved protein sequences were aligned using webPRANK (Löytynoja & Goldman, 2010) and a maximum likelihood phylogeny was built using MEGA X (Kumar *et al.*, 2018).
